# A Combinatorial PCR Method for Efficient, Selective Oligo Retrieval from Complex Oligo Pools

**DOI:** 10.1101/2021.08.25.457714

**Authors:** Claris Winston, Lee Organick, Luis Ceze, Karin Strauss, Yuan-Jyue Chen

## Abstract

With the rapidly decreasing cost of array-based oligo synthesis, large-scale oligo pools offer significant benefits for advanced applications, including gene synthesis, CRISPR-based gene editing, and DNA data storage. Selectively retrieving specific oligos from these complex pools traditionally uses Polymerase Chain Reaction (PCR), in which any selected oligos are exponentially amplified to quickly outnumber non-selected ones. In this case, the number of orthogonal PCR primers is limited due to interactions between them. This lack of specificity presents a serious challenge, particularly for DNA data storage, where the size of an oligo pool (i.e., a DNA database) is orders of magnitude larger than it is for other applications. Although a nested file address system was recently developed to increase the number of accessible files for DNA storage, it requires a more complicated lab protocol and more expensive reagents to achieve high specificity. Instead, we developed a new combinatorial PCR method that outperforms prior work without compromising the fidelity of retrieved material or complicating wet lab processes. Our method quadratically increases the number of accessible oligos while maintaining high specificity. In experiments, we accessed three arbitrarily chosen files from a DNA prototype database that contained 81 different files. Initially comprising only 1% of the original database, the selected files were enriched to over 99.9% using our combinatorial primer method. Our method thus provides a viable path for scaling up DNA data storage systems and has broader utility whenever scientists need access to a specific target oligo and can design their own primer regions.

## Introduction

De novo synthesis of nucleic acids plays a key role in understanding and engineering biology and in enabling development of novel applications in nanotechnology and DNA data storage.^1,2^ The use of synthetic DNA has been significantly aided by the development of large-scale, array-based DNA synthesis.^3^ Compared to traditional column-based oligo synthesis,^4–6^ arrays make multiple oligos in parallel, and all synthesized oligos are cleaved and harvested as one ‘oligo pool.’^7–10^ This new development has dramatically decreased the cost of oligonucleotide synthesis, providing the foundation for large-scale gene synthesis and enabling the large-scale encoding of information in DNA for long-term data storage.^11,12^

Although array-based oligo pools are cheap and rapidly becoming cheaper^3^, they pose a challenging new question: as thousands to millions of oligos are made together in a single oligo pool, how can we selectively retrieve one or more specific oligos (e.g., a gene fragment or a data file encoded in DNA)?

This problem has traditionally been solved by flanking target sequence(s) with a unique pair of primers (**Figure 1a**). Target sequences (or “files”) are retrieved by PCR amplification using corresponding primers. However, with thousands of segments, each with one unique forward and one unique reverse primer,^13,14^ ordering thousands of primers can become cost prohibitive. This problem is particularly concerning for DNA data storage, where millions to billions of DNA files could be stored together, mandating the development of an efficient, uncomplicated, and inexpensive random access method to retrieve a given target file.^14–16^

**Figure 1.**
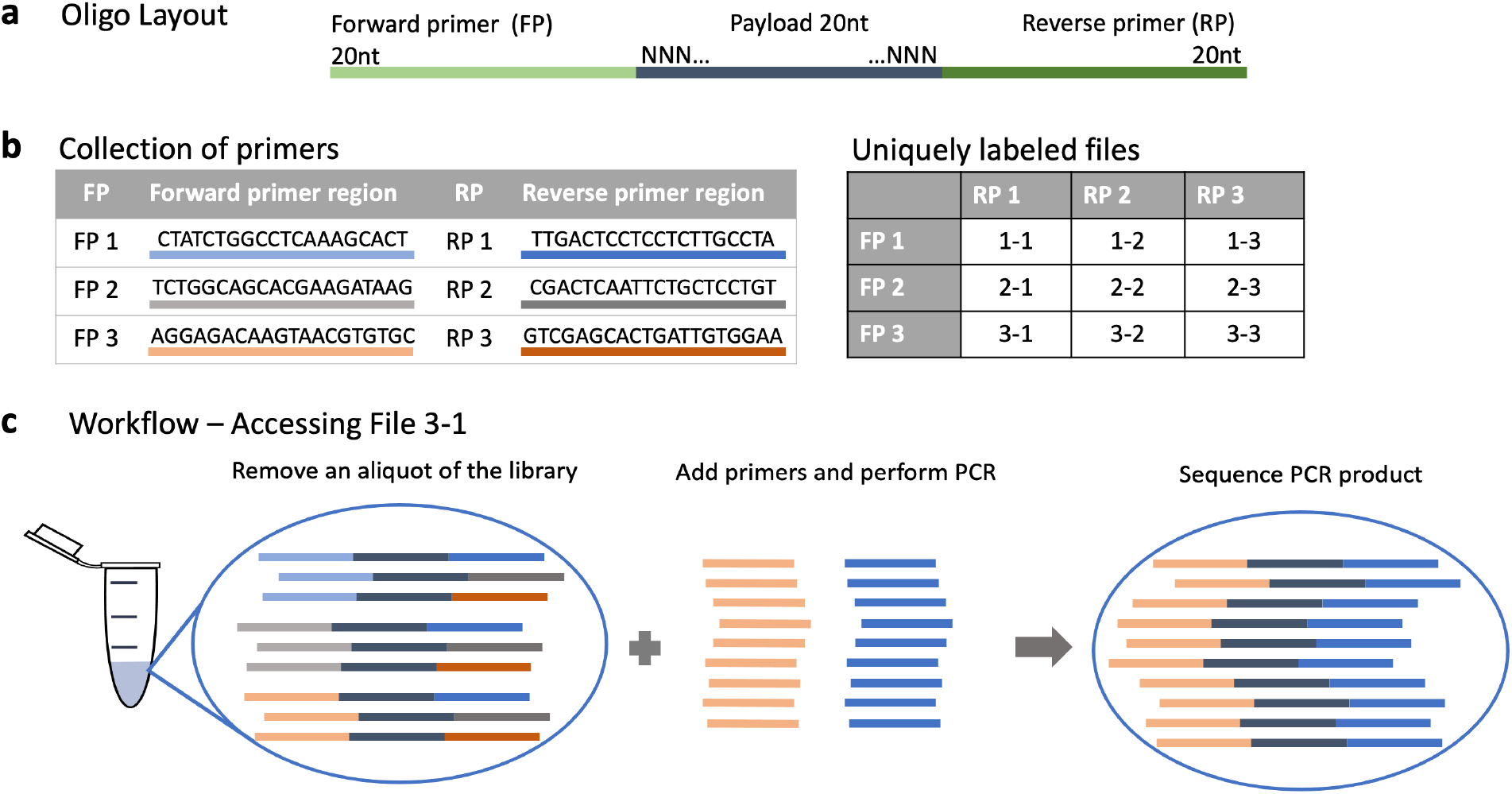
Overview of the primer method and protocol. (a) Oligo layout. The DNA of interest (data, genomic material, etc.) is stored in the middle portion of the oligo, flanked by forward primer (FP) and reverse primer (RP) binding regions. In this work, the FP is 20 nucleotides (nt) long, the middle section is 20 nt, and the RP is 20 nt, for a total strand length of 60 nt. (b) Left: A sequence table of six primer sequences used in this work. Right: Primers can be re-used in different files, but each file is labeled with a unique pair of primers. For example, File 1-1 uses FP1 and RP1. File 2-1 uses FP2 and RP1. **Supplemental Section 1** shows a complete list of all sequences used in this work. (c) A schematic showing the standard PCR where two primers retrieve the corresponding target file.

In a recently presented alternative hierarchical addressing method with a universal reverse primer and two nested forward primers,^15^ ten primers uniquely identified and retrieved 72 files; contrast this with the traditional “one unique primer at each end” method, where ten primers can uniquely identify only five files (**Figure 2**). However, while the hierarchical primer design is the most primer efficient method to date, there are several key drawbacks.

**Figure 2.**
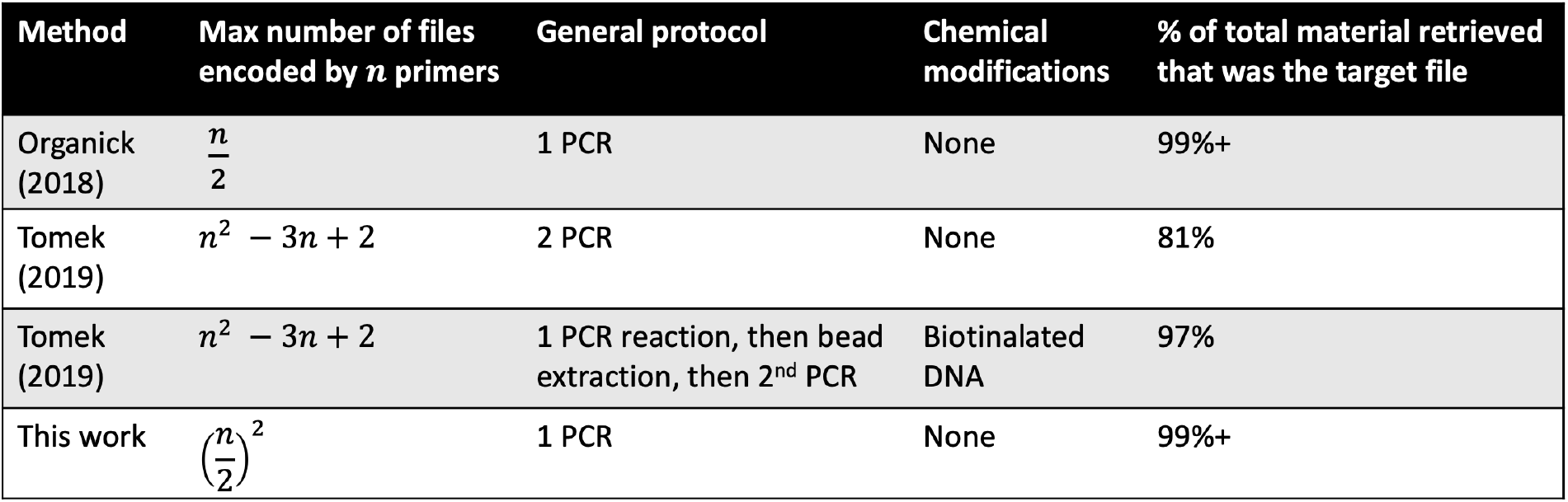
A comparison of various file addressing methods. Shown are the scalability of the number of primers needed, the general protocol complexity of retrieving the target file, whether any chemical modification is necessary at any stage of the wet lab protocol used to recover the target file, and retrieval efficiency.

First, the hierarchical addressing scheme does not retrieve target strands with nearly as high fidelity due to the use of a universal reverse primer region to amplify every sequence, with retrieval fidelity of either 81% or 97% depending on the exact methods used compared to this work’s fidelity of 99.9%+. Returning only target files and sequences with high fidelity is vital for several reasons. Some applications cannot tolerate errant sequences (i.e., targeted genome modification). Furthermore, sequencing resources are wasted on non-target sequences. In the context of DNA data storage, where each data file is encoded in thousands to many million sequences of DNA, having only 81% of returned material be the target file^15^ can require costly amounts of extra sequencing. Second, to achieve the 97% fidelity, the wet lab retrieval methods are more involved and require additional time and reagents.^15^ While this may not be prohibitive to some applications, others may find the extra steps prohibitively difficult to automate, or too time intensive to make the trade off advantageous. Third, by requiring three addresses per DNA strand rather than the traditional two, some applications may find that their DNA strand is prohibitively long or expensive to synthesize. While the hierarchical address method may be a wonderful solution for some applications who prioritize the fewest number of primer sequences, some users may find one of these drawbacks prohibitive for their specific application but still wish to minimize the number of primer sequences necessary.

This paper introduces a combinatorial PCR method that outperforms the traditional PCR random access method (i.e., one unique forward and reverse primer per file) without compromising the fidelity of retrieved material or complicating protocols. We design a collection of forward and of reverse primer sequences that are paired combinatorially to allow every unique forward-reverse primer combination. In this design, ten total primer sequences (five forward and five reverse primers) can uniquely identify 25 files (**Figure 1a, 1b**), and the target file is simply accessed with standard PCR (**Figure 1c**). While this method requires more primers than the Tomek et al. approach,^15^ it is a more efficient, inexpensive protocol: it requires no bead extraction or DNA modification and yields much less unwanted amplification (<0.1%). Figure 2 compares these two methods.

## Results

### Experimental Sequencing Retrieval Results

We tested our method in a prototype oligo pool in which 9 forward and 9 reverse primers were used to specify 81 unique files. This system was at “full capacity” since all 81 possible pairings of forward and reverse primers were used to specify the 81 unique files. To simulate a highly complex pool, each file was ordered with completely random bases between primer regions. The diversity of the random bases (20 random bases yielding 4^20^ possible sequences) was sufficiently large to make it unlikely that any two retrieved DNA strands were the same.

We performed PCR to amplify three arbitrarily chosen files from the pool. The amplified files were 1-1 (specified by FP1 and RP1), 2-2 (specified by FP2 and RP2), and 3-3 (specified by FP3 and RP3). We ran five separate PCR reactions to amplify each file (we ran duplicates for files 2-2 and 3-3) from the pool of 81 files. Each of the resulting five samples was sequenced to reveal the amount of target and non-target oligos in the amplified material. If our retrieval method performed perfectly, we would expect to see: virtually all sequencing reads correspond to the target file, a small proportion of reads corresponding to files that share one primer with the target file and were linearly amplified, and a very small proportion of sequences from non-target files as a result of non-amplified DNA present in the original subsample of the pool used at the start of the PCR reaction. As **Figure 3** shows, the percentage of the target file in all 5 experiments exceeded 99.9%, indicating near-perfect retrieval efficiency. Importantly, there was a negligible variation of only 0.062% observed between all five experiments’ retrieval efficiencies, demonstrating this primer method’s consistent high fidelity retrieval.

**Figure 3.**
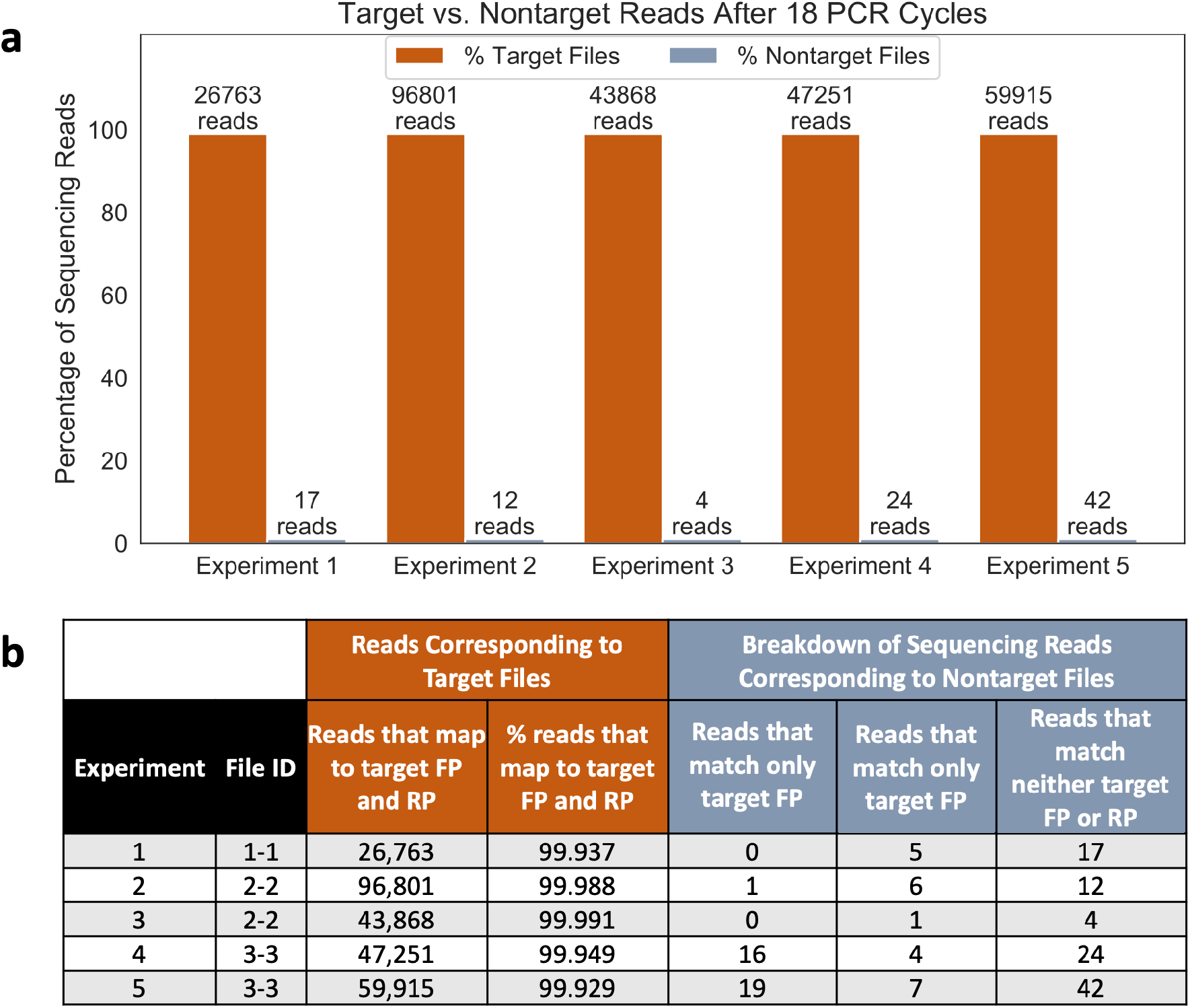
Analyzing sequencing data. (a) Percentage of sequencing reads that belong to target files (orange bars) and non-target files (blue bars). (b) More detail about the sequencing reads for each experiment. Sequencing reads corresponding to a target file are defined as reads mapped to both target FP and RP (in orange). Reads corresponding to non-target files (in blue) have either only one or zero primers matching the target file primers.

We investigated whether the recovered non-target sequences had been linearly amplified due to sharing either a forward or reverse primer and whether recovered non-target strands were arbitrary strands left over from subsampling the initial pool; **Figure 3b** shows a condensed table of these results. Compared to tens of thousands of sequencing reads corresponding to our target file, both linearly amplified sequences and non-target sequences showed extremely low retrieval (low 10s of sequencing reads). Linearly amplified sequences showed even fewer reads than non-target strands that did not match forward or reverse primers of our target sequences. This could be due to imperfect PCR efficiency, non-specific carry-over from the oligo pool, and the fact that most post-PCR linearly amplified sequences are single-stranded, and single-stranded oligos are filtered out during the sequencing library preparation (**Supplementary Section 2**).

### Simulation of Retrieval Efficiency from a Complex Oligo Pool

We further modeled this system to understand how the result might change when the pool contains many more files. In our experiment, we implemented and analyzed a system with 9 forward and 9 reverse primers to specify 81 files. Here, we generalize these results to a oligo pool where there are *a* forward primers and *b* reverse primers. The maximum number *a*^2^ of addresses is achieved when *a* = *b* because then we can create *a · b* primer pairs to specify sequences. The retrieval efficiency *χ* denotes the fraction of the molecule number of a target oligo over the total number of molecules in a pool after PCR:

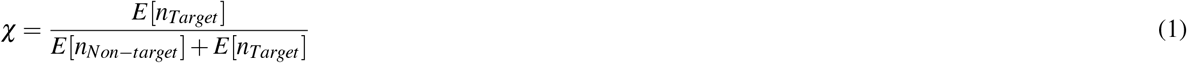

Next, we examine what effect the number of PCR cycles and number of initial molecules has on the expected final number of molecules. We start with *n*_0_ molecules in a DNA pool and assume that all files are the same size, and each file has *n*_0_/*a*^2^ molecules. In each PCR cycle, each molecule is amplified with the probability *p*, where 0 *≤ p ≤* 1. PCR amplification is a branching process.^17,18^

The expected molecular number of a target file after *i* cycle(s) of PCR is:

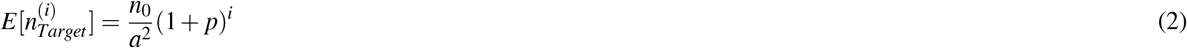

The expected molecular number of other non-target files after *i* cycle(s) of PCR is:

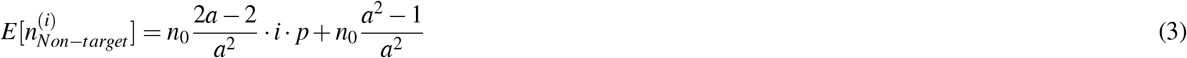

The first and second terms of Equation (3) correspond to files that are linearly amplified and not amplified, respectively.

The random access efficiency of a target file *χ* is calculated as follows:

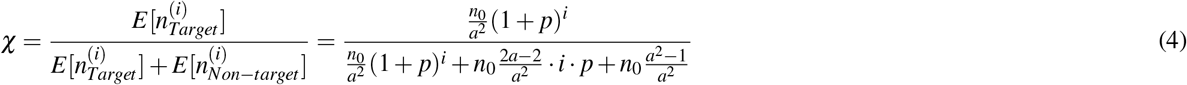

To see how random access efficiency *χ* changes as the number of files in a pool increases, we plot *χ* against the number of files (**Supplementary Figure 1**). Random access efficiency decreases as more files are used in a database. Despite the linear amplification that occurs, *χ* can dramatically improve when more PCR cycles are used. Our experiment used 18 cycles to amplify a target file in our 81-file system. We modeled this system visually using more files (**Supplementary Figure 1**).

If we used the same number of cycles in much larger systems, e.g., 1,000,000 files, the proportion of a target files after 18 cycles would be less than 0.2, which indicates very low efficiency. We could circumvent this by using more PCR cycles. For example, if we increase the number of PCR cycles to 25, the proportion of target file can achieve *∼*0.7. However, doing so increases the risk of primer dimers or of a primer running out and biasing the sample. Another option could be a two-stage PCR, where additional primer is added between the stages; this would let us increase large-system efficiency by adding more PCR cycles without the high risk of primer dimers or severe biasing.

## Discussion

We developed and evaluated a new combinatorial PCR method to selectively retrieve oligos from a complex oligo pool. Ours is a more efficient method of creating files than traditional primer design, which does not use primers combinatorially, and we experimentally demonstrated that it exclusively retrieved target sequences at a high efficiency of over 99.9%. Furthermore, the combinatorial implementation we evaluated did not rely on expensive DNA modification or additional reagents;^15,16,19^ instead, it simply employed standard PCR for less-expensive retrieval requiring fewer primers and was just as quick and easy to perform as traditional PCR.

These properties – the number of primers, protocol cost, speed, and simplicity – are important considerations to balance for any primer design. For example, in the rapidly growing field of DNA data storage, many files are stored in the same pool and are preserved for many years before they are accessed, if ever. In this scenario, the number of primers used is a negligible cost compared to the time and financial cost of a more complex access protocol, especially compared to the cost of sequencing resources wasted on non-target sequences. However, in a synthetic gene context the sequencing cost might be negligible, but if the primer design scheme requires the strand to be a longer length the cost of the strand may become too great, or a retrieval protocol more time and resource intensive than a standard PCR may be undesirable.

Our computational model shows the possibility of scaling to DNA databases with up to millions of files while maintaining high retrieval efficiency. It is worth noting that to achieve high retrieval efficiency in a very large library (for example, a million files), it requires to use more cycles of PCR and primers. If the formation of primer dimers is a concern, a two-stage PCR, where additional primer is added between the stages, could be used to circumvent such issue. In applications such as DNA data storage, where each file might contain many sequences, the scalability, simplicity, and high efficiency of our combinatorial PCR method provides a viable path towards building a practical DNA storage system. We also stress that this PCR primer method is potentially useful in any context where scientists might need to access a specific target oligo and have the flexibility to design their own primer regions.

## Methods

### Primer Design

All primer region sequences are shown in **Supplemental Section 1**. Primer sequences were designed using the sequence design method in large-scale DNA data storage.^20^ In brief, primer sequences were optimized with the following criteria: (1) GC content between 45% and 55%, (2) absence of long homopolymers (i.e., AAAA, TTTT, GGG, and CCC), (3) absence of more than 4-bp self-sequence complementarity, (4) absence of more than 10-bp inter-sequence complementarity, (5) Hamming distance *≤*6, and (6) avoidance of secondary structure. The basic sequence alignment program BLAST was used to screen out primers with long stretches of similar sequences.

### DNA Reagents

All oligos were synthesized by Integrated DNA Technologies. DNA templates (mimicking DNA files) and primers were purchased as desalted, unpurified DNA. The 20N payload was comprised of IDT’s standard “N” composition (where A/T/G/C are randomly added). All sequences are shown in **Supplemental Section 1**. All DNA was resuspended to 100 uM in 1X TE buffer (pH7.5). KAPA HIFI polymerase was purchased from Kapa Biosystems. Ligation reagents were part of Illumina’s TruSeq Nano kit.

### PCR: Amplifying Files

In a 20 uL PCR reaction, 1 uL of 10 nM of the ordered IDT ssDNA pool was mixed with 1 uL of 10 uM of the forward and 10 uM of the reverse primer, 10 uL of 2X KAPA HIFI enzyme mix, and 8 uL of molecular biograde water. The reaction followed a thermal protocol: (1) 95 ^*º*^C for 3 min, (2) 98 ^*º*^C for 20 s, (3) 62 ^*º*^C for 20 s, and (4) 72 ^*º*^C for 15 s. The size of PCR products was confirmed using a Qiaxcel fragment analyzer.

### Quantitative PCR (qPCR)

In each 20 uL qPCR reaction, 1 uL of the ssDNA pool was mixed with 1 uL of the respective forward and reverse primer mix, 1 uL of 20X EvaGreen dye, 10 uL of 2X KAPA HIFI enzyme mix, and 7 uL of molecular water. The reaction followed a thermal protocol: (1) 95 ^*º*^C for 3 min, and 39 cycles of (2) 98 ^*º*^C for 20 s, (3) 62 ^*º*^C for 20 s, and (4) 72 ^*º*^C for 15 s.

### Sample Preparation for Sequencing

PCR products were ligated to Illumina adapters by following a modified version of a combination of the Illumina TrueSeq Nano DNA Library Prep and TruSeq ChiP Sample preparation protocol using the TruSeq Nano kit. Samples were converted to blunt ends by following the “End Repair” step in the Illumina TruSeq Nano kit, then purified with AMPure XP beads using the TruSeq ChiP protocol. An ‘A’ base was added to the 3’ end of the DNA fragments with the TruSeq Nano’s A-tailing protocol, followed by ligation to the Illumina adapters with the TruSeq Nano protocol. The samples were then cleaned with Illumina sample purification beads and enriched using PCR to generate sufficient products for sequencing. The sample size was confirmed using a Qiaxcel Bioanalyzer. A step-by-step ligation protocol is shown in **Supplemental Section 2**. Immediately before sequencing, the sample concentration was measured using qPCR. The sample was then prepared for sequencing following the *NextSeq System Denature and Dilute Libraries Guide*. The sample was loaded into a sequencer at 1.3 pM with a 10 *−* 20% PhiX spike in as a control.

### Processing Sequencing Data

After the PCR results were sequenced, the *.fasta* files were processed using a BLAST command, which can be found in file *Sequencing Analysis*.*ipynb* in the GitHub repository given in the code disclosure below. The BLAST command looks for matches of primer sequences in the original sequences and results in a *.txt* file, with the rows containing the matching sequences along with alignment values. The BLAST command we used had a maximum of 1000000 target sequences found (max target seqs), a reward of 1 for a nucleotide match (reward), and a penalty of -2 for a nucleotide mismatch (penalty).^21^ We next ran a Python script (*Sequencing Analysis*.*ipynb*, found in the Github repository) that first filtered the *.txt* files output from the BLAST command so they included only sequences no longer than 85 nucleotides and had an alignment region of at least 15. This reduced noise in the sequencing analysis. Finally, we used the script to count the number of copies of each file. Each sequence should have appeared two times in the *.txt* file, once when the forward primer was aligned with that sequence, and once when the reverse primer was aligned with that sequence. The Python code looks for these repeated instances of the same sequence in the output file and maps each sequence to the forward and reverse primer pair (this pair represents a file) corresponding to it. So, by counting the number of sequences that mapped to each possible primer pair, we could count the number of copies of each file in the pool since each file is specified by a unique primer pair.

## Data Availability Statement

Data are available at https://github.com/uwmisl/combinatorial_primer_project.

## Code Availability Statement

Code used to perform analyses presented in the paper is available at https://github.com/uwmisl/combinatorial_primer_project.

## Acknowledgements

The authors are grateful to Sandy Kaplan for editing and providing feedback on the manuscript, and to Jeff Nivala for providing feedback on the manuscript.

Funding provided by DARPA Molecular Informatics Program and Microsoft.

## Author Contributions Statement

C.W. performed wet lab experiments, and C.W., L.O., and Y.J.C analyzed the data and wrote the manuscript. Y.J.C., L.C., and K.S. directed and supervised the work.

