## Supplemental Information for "A Combinatorial PCR Method for Efficient, Selective Oligo Retrieval from Complex Oligo Pools"

Supplementary Material

Winston *et al.*

### Contents


**1 Performing and Analyzing qPCR** **3**

1.1 Primer Sequences . . . 3

1.2 81 File Sequences . . . 3

**2 Ligation Protocol** **7**

**3 Modeling Large Scale Pools** **11**

### 1 Performing and Analyzing qPCR

In the following two subsections, we have listed the the primer region sequences and the sequences for the files used in this work. The N denotes an equal chance of being an A, G, T, or C. It is important to note that all sequences listed are the raw sequences ordered from IDT, listed from 5' to 3'.

#### 1.1 Primer Sequences

This table has the 9 forward primer and 9 reverse primer sequences that appear in the files. If the primer id starts with 'FP', that indicates the sequence corresponds to a forward primer, and if it starts with 'RP', that indicates the sequence corresponds to a reverse primer. Below are the sequences and their primer IDs:

| ID | Sequence |
|---|---|
| FP1 | CTATCTGGCCTCAAAGCACT |
| FP2 | TCTGGCAGCACGAAGATAAG |
| FP3 | AGGAGACAAGTAACGTGTGC |
| FP4 | TAAGAAAGGCATCGTCACCG |
| FP5 | GTCACGTCACCAGGTAACAA |
| FP6 | TTTCCGATAGTTGAGGCAGG |
| FP7 | TGGTGAAACTACCGACTTCC |
| FP8 | AGTTGGTGACTATCCGTCCT |
| FP9 | CACATAGGCAAAGCGGAGTA |
| RP1 | TTGACTCCTCCTCTTGCCTA |
| RP2 | CGACTCAATTCTGCTCCTGT |
| RP3 | GTCGAGCACTGATTGTGGAA |
| RP4 | GTACTGCTCGGCCACTTATT |
| RP5 | GCGTTGTCTCTAGCGAAAGA |
| RP6 | TCATTTCTCCGACAGGCTTG |
| RP7 | CCTCTCTTCGCGTTGATCTT |
| RP8 | GCAAGACAATAGGCTTCGGT |
| RP9 | GCAAGACAATAGGCTTCGGT |

Note that to perform PCR on a file with these sequences, one should order the order the reverse complement of the RP sequence given here.

#### 1.2 81 File Sequences

These are the full sequences to all the 81 files along with their ids, again listed from 5' to 3'. The naming scheme for the files are as follows:
[number associated with forward primer]-[number associated with reverse primer]
For example, the file id '4-5' indicates that the file is specified by FP4 and RP5.

Below are all 81 sequences that correspond to each of these files:

| ID | Sequence |
|---|---|
| 1-1 | CTATCTGGCCTCAAAGCACTNNNNNNNNNNNNNNNNNNNNTTGACTCCTCCTCTTGCCTA |
| 1-2 | CTATCTGGCCTCAAAGCACTNNNNNNNNNNNNNNNNNNNNCGACTCAATTCTGCTCCTGT |
| 1-3 | CTATCTGGCCTCAAAGCACTNNNNNNNNNNNNNNNNNNNNGTCGAGCACTGATTGTGGAA |
| 1-4 | CTATCTGGCCTCAAAGCACTNNNNNNNNNNNNNNNNNNNNGTACTGCTCGGCCACTTATT |
| 1-5 | CTATCTGGCCTCAAAGCACTNNNNNNNNNNNNNNNNNNNNGCGTTGTCTCTAGCGAAAGA |
| 1-6 | CTATCTGGCCTCAAAGCACTNNNNNNNNNNNNNNNNNNNNTCATTTCTCCGACAGGCTTG |
| 1-7 | CTATCTGGCCTCAAAGCACTNNNNNNNNNNNNNNNNNNNNCCTCTCTTCGCGTTGATCTT |
| 1-8 | CTATCTGGCCTCAAAGCACTNNNNNNNNNNNNNNNNNNNNGCAAGACAATAGGCTTCGGT |
| 1-9 | CTATCTGGCCTCAAAGCACTNNNNNNNNNNNNNNNNNNNNCGCTGCTGGTAATTTAACCG |
| 2-1 | TCTGGCAGCACGAAGATAAGNNNNNNNNNNNNNNNNNNNNTTGACTCCTCCTCTTGCCTA |
| 2-2 | TCTGGCAGCACGAAGATAAGNNNNNNNNNNNNNNNNNNNNCGACTCAATTCTGCTCCTGT |
| 2-3 | TCTGGCAGCACGAAGATAAGNNNNNNNNNNNNNNNNNNNNGTCGAGCACTGATTGTGGAA |
| 2-4 | TCTGGCAGCACGAAGATAAGNNNNNNNNNNNNNNNNNNNNGTACTGCTCGGCCACTTATT |
| 2-5 | TCTGGCAGCACGAAGATAAGNNNNNNNNNNNNNNNNNNNNGCGTTGTCTCTAGCGAAAGA |
| 2-6 | TCTGGCAGCACGAAGATAAGNNNNNNNNNNNNNNNNNNNNTCATTTCTCCGACAGGCTTG |
| 2-7 | TCTGGCAGCACGAAGATAAGNNNNNNNNNNNNNNNNNNNNCCTCTCTTCGCGTTGATCTT |
| 2-8 | TCTGGCAGCACGAAGATAAGNNNNNNNNNNNNNNNNNNNNGCAAGACAATAGGCTTCGGT |
| 2-9 | TCTGGCAGCACGAAGATAAGNNNNNNNNNNNNNNNNNNNNCGCTGCTGGTAATTTAACCG |
| 3-1 | AGGAGACAAGTAACGTGTGCNNNNNNNNNNNNNNNNNNNNTTGACTCCTCCTCTTGCCTA |
| 3-2 | AGGAGACAAGTAACGTGTGCNNNNNNNNNNNNNNNNNNNNCGACTCAATTCTGCTCCTGT |
| 3-3 | AGGAGACAAGTAACGTGTGCNNNNNNNNNNNNNNNNNNNNGTCGAGCACTGATTGTGGAA |
| 3-4 | AGGAGACAAGTAACGTGTGCNNNNNNNNNNNNNNNNNNNNGTACTGCTCGGCCACTTATT |
| 3-5 | AGGAGACAAGTAACGTGTGCNNNNNNNNNNNNNNNNNNNNGCGTTGTCTCTAGCGAAAGA |
| 3-6 | AGGAGACAAGTAACGTGTGCNNNNNNNNNNNNNNNNNNNNTCATTTCTCCGACAGGCTTG |
| 3-7 | AGGAGACAAGTAACGTGTGCNNNNNNNNNNNNNNNNNNNNCCTCTCTTCGCGTTGATCTT |
| 3-8 | AGGAGACAAGTAACGTGTGCNNNNNNNNNNNNNNNNNNNNGCAAGACAATAGGCTTCGGT |
| 3-9 | AGGAGACAAGTAACGTGTGCNNNNNNNNNNNNNNNNNNNNCGCTGCTGGTAATTTAACCG |
| 4-1 | TAAGAAAGGCATCGTCACCGNNNNNNNNNNNNNNNNNNNNTTGACTCCTCCTCTTGCCTA |
| 4-2 | TAAGAAAGGCATCGTCACCGNNNNNNNNNNNNNNNNNNNNCGACTCAATTCTGCTCCTGT |
| 4-3 | TAAGAAAGGCATCGTCACCGNNNNNNNNNNNNNNNNNNNNGTCGAGCACTGATTGTGGAA |
| 4-4 | TAAGAAAGGCATCGTCACCGNNNNNNNNNNNNNNNNNNNNGTACTGCTCGGCCACTTATT |
| 4-5 | TAAGAAAGGCATCGTCACCGNNNNNNNNNNNNNNNNNNNNGCGTTGTCTCTAGCGAAAGA |
| 4-6 | TAAGAAAGGCATCGTCACCGNNNNNNNNNNNNNNNNNNNNTCATTTCTCCGACAGGCTTG |
| 4-7 | TAAGAAAGGCATCGTCACCGNNNNNNNNNNNNNNNNNNNNCCTCTCTTCGCGTTGATCTT |
| 4-8 | TAAGAAAGGCATCGTCACCGNNNNNNNNNNNNNNNNNNNNGCAAGACAATAGGCTTCGGT |
| 4-9 | TAAGAAAGGCATCGTCACCGNNNNNNNNNNNNNNNNNNNNCGCTGCTGGTAATTTAACCG |

| | |
|---|---|
| 5-1 | GTCACGTCACCAGGTAACAANNNNNNNNNNNNNNNNNNNNTTGACTCCTCCTCTTGCCTA |
| 5-2 | GTCACGTCACCAGGTAACAANNNNNNNNNNNNNNNNNNNNCGACTCAATTCTGCTCCTGT |
| 5-3 | GTCACGTCACCAGGTAACAANNNNNNNNNNNNNNNNNNNNGTCGAGCACTGATTGTGGAA |
| 5-4 | GTCACGTCACCAGGTAACAANNNNNNNNNNNNNNNNNNNNGTACTGCTCGGCCACTTATT |
| 5-5 | GTCACGTCACCAGGTAACAANNNNNNNNNNNNNNNNNNNNGCGTTGTCTCTAGCGAAAGA |
| 5-6 | GTCACGTCACCAGGTAACAANNNNNNNNNNNNNNNNNNNNTCATTTCTCCGACAGGCTTG |
| 5-7 | GTCACGTCACCAGGTAACAANNNNNNNNNNNNNNNNNNNNCCTCTCTTCGCGTTGATCTT |
| 5-8 | GTCACGTCACCAGGTAACAANNNNNNNNNNNNNNNNNNNNGCAAGACAATAGGCTTCGGT |
| 5-9 | GTCACGTCACCAGGTAACAANNNNNNNNNNNNNNNNNNNNCGCTGCTGGTAATTTAACCG |
| 6-1 | TTTCCGATAGTTGAGGCAGGNNNNNNNNNNNNNNNNNNNNTTGACTCCTCCTCTTGCCTA |
| 6-2 | TTTCCGATAGTTGAGGCAGGNNNNNNNNNNNNNNNNNNNNCGACTCAATTCTGCTCCTGT |
| 6-3 | TTTCCGATAGTTGAGGCAGGNNNNNNNNNNNNNNNNNNNNGTCGAGCACTGATTGTGGAA |
| 6-4 | TTTCCGATAGTTGAGGCAGGNNNNNNNNNNNNNNNNNNNNGTACTGCTCGGCCACTTATT |
| 6-5 | TTTCCGATAGTTGAGGCAGGNNNNNNNNNNNNNNNNNNNNGCGTTGTCTCTAGCGAAAGA |
| 6-6 | TTTCCGATAGTTGAGGCAGGNNNNNNNNNNNNNNNNNNNNTCATTTCTCCGACAGGCTTG |
| 6-7 | TTTCCGATAGTTGAGGCAGGNNNNNNNNNNNNNNNNNNNNCCTCTCTTCGCGTTGATCTT |
| 6-8 | TTTCCGATAGTTGAGGCAGGNNNNNNNNNNNNNNNNNNNNGCAAGACAATAGGCTTCGGT |
| 6-9 | TTTCCGATAGTTGAGGCAGGNNNNNNNNNNNNNNNNNNNNCGCTGCTGGTAATTTAACCG |
| 7-1 | TGGTGAAACTACCGACTTCCNNNNNNNNNNNNNNNNNNNNTTGACTCCTCCTCTTGCCTA |
| 7-2 | TGGTGAAACTACCGACTTCCNNNNNNNNNNNNNNNNNNNNCGACTCAATTCTGCTCCTGT |
| 7-3 | TGGTGAAACTACCGACTTCCNNNNNNNNNNNNNNNNNNNNGTCGAGCACTGATTGTGGAA |
| 7-4 | TGGTGAAACTACCGACTTCCNNNNNNNNNNNNNNNNNNNNGTACTGCTCGGCCACTTATT |
| 7-5 | TGGTGAAACTACCGACTTCCNNNNNNNNNNNNNNNNNNNNGCGTTGTCTCTAGCGAAAGA |
| 7-6 | TGGTGAAACTACCGACTTCCNNNNNNNNNNNNNNNNNNNNTCATTTCTCCGACAGGCTTG |
| 7-7 | TGGTGAAACTACCGACTTCCNNNNNNNNNNNNNNNNNNNNCCTCTCTTCGCGTTGATCTT |
| 7-8 | TGGTGAAACTACCGACTTCCNNNNNNNNNNNNNNNNNNNNGCAAGACAATAGGCTTCGGT |
| 7-9 | TGGTGAAACTACCGACTTCCNNNNNNNNNNNNNNNNNNNNCGCTGCTGGTAATTTAACCG |
| 8-1 | AGTTGGTGACTATCCGTCCTNNNNNNNNNNNNNNNNNNNNTTGACTCCTCCTCTTGCCTA |
| 8-2 | AGTTGGTGACTATCCGTCCTNNNNNNNNNNNNNNNNNNNNCGACTCAATTCTGCTCCTGT |
| 8-3 | AGTTGGTGACTATCCGTCCTNNNNNNNNNNNNNNNNNNNNGTCGAGCACTGATTGTGGAA |
| 8-4 | AGTTGGTGACTATCCGTCCTNNNNNNNNNNNNNNNNNNNNGTACTGCTCGGCCACTTATT |
| 8-5 | AGTTGGTGACTATCCGTCCTNNNNNNNNNNNNNNNNNNNNGCGTTGTCTCTAGCGAAAGA |
| 8-6 | AGTTGGTGACTATCCGTCCTNNNNNNNNNNNNNNNNNNNNTCATTTCTCCGACAGGCTTG |
| 8-7 | AGTTGGTGACTATCCGTCCTNNNNNNNNNNNNNNNNNNNNCCTCTCTTCGCGTTGATCTT |
| 8-8 | AGTTGGTGACTATCCGTCCTNNNNNNNNNNNNNNNNNNNNGCAAGACAATAGGCTTCGGT |
| 8-9 | AGTTGGTGACTATCCGTCCTNNNNNNNNNNNNNNNNNNNNCGCTGCTGGTAATTTAACCG |

| | |
|---|---|
| 9-1 | CACATAGGCAAAGCGGAGTANNNNNNNNNNNNNNNNNNNNTTGACTCCTCCTCTTGCCTA |
| 9-2 | CACATAGGCAAAGCGGAGTANNNNNNNNNNNNNNNNNNNNCGACTCAATTCTGCTCCTGT |
| 9-3 | CACATAGGCAAAGCGGAGTANNNNNNNNNNNNNNNNNNNNGTCGAGCACTGATTGTGGAA |
| 9-4 | CACATAGGCAAAGCGGAGTANNNNNNNNNNNNNNNNNNNNGTACTGCTCGGCCACTTATT |
| 9-5 | CACATAGGCAAAGCGGAGTANNNNNNNNNNNNNNNNNNNNGCGTTGTCTCTAGCGAAAGA |
| 9-6 | CACATAGGCAAAGCGGAGTANNNNNNNNNNNNNNNNNNNNTCATTTCTCCGACAGGCTTG |
| 9-7 | CACATAGGCAAAGCGGAGTANNNNNNNNNNNNNNNNNNNNCCTCTCTTCGCGTTGATCTT |
| 9-8 | CACATAGGCAAAGCGGAGTANNNNNNNNNNNNNNNNNNNNGCAAGACAATAGGCTTCGGT |
| 9-9 | CACATAGGCAAAGCGGAGTANNNNNNNNNNNNNNNNNNNNCGCTGCTGGTAATTTAACCG |

#### 2 Ligation Protocol

The ligation protocol used is a combination of the Illumina TruSeq ChIP Sample Preparation protocol and the Illumina TruSeq Nano Library Prep protocol, using the Illumina TruSeq Nano reagents. The following are the step-by-step directions used in this work.

1. Add 40 $\mu$L ERP2 to each well with 60 $\mu$L of PCR product
2. Pipette up and down to mix
3. Place on the thermal cycler and run the ERP program (30 min at 30C) then place on ice. Each well contains 100 $\mu$L.
4. Add 160 $\mu$L well-mixed AMPure XP Beads to each well of the PCR plate containing 100 $\mu$L End Repair Mix.
5. Gently pipette the entire volume up and down 10 times to mix thoroughly.
6. Incubate the PCR plate at room temperature for 15 minutes.
7. Place the PCR plate on a magnetic stand at room temperature for 15 minutes or until the liquid is clear.
8. Using a 200 $\mu$L single channel or multichannel pipette set to 127.5 $\mu$L, remove and discard 127.5 $\mu$L of the supernatant from each well of the PCR plate.
9. Repeat step 8 one time.

   NOTE- Leave the PCR plate on the magnetic stand while performing the following 80% EtOH wash steps (10-12).
10. With the PCR plate on the magnetic stand, add 200 $\mu$L freshly prepared 80% EtOH to each well without disturbing the beads.
11. Incubate the PCR plate at room temperature for 30 seconds, and then remove and discard all of the supernatant from each well. Take care not to disturb the beads.
12. Repeat steps 10 and 11 one time for a total of two 80% EtOH washes.
13. Let the PCR plate stand at room temperature for 15 minutes to dry, and then remove the plate from the magnetic stand.
14. Resuspend the dried pellet in each well with 17.5 $\mu$L Resuspension Buffer (RSB). Gently pipette the entire volume up and down 10 times to mix thoroughly.
15. Incubate the PCR plate at room temperature for 2 minutes.

16. Place the PCR plate on the magnetic stand at room temperature for 5 minutes or until the liquid is clear.
17. Transfer 15 $\mu$L of the clear supernatant from each well of the PCR plate to the corresponding well of a new 96-well 0.3 ml PCR plate.
18. Add 15 $\mu$L of A Tailing Ligase (ATL) to the 15 $\mu$L of supernatant from previous step.
19. Briefly spin down on a standard, small benchtop centrifuge.
20. Place on the thermal cycler and run the ATAIL70 program listed below. Each well contains 30 $\mu$L.
    (a) Choose the preheat lid option and set to 100°C
        i. 37C for 30 minutes
        ii. 70C for 5 minutes
        iii. 4C for 5 minutes
21. Add the following reagents from the Illumina TruSeq Nano ligation kit IN ORDER:
    (a) RSB (2.5 $\mu$L)
    (b) LIG2 (2.5 $\mu$L)
    (c) DNA adapter (2.5 $\mu$L)
22. Pipette up and down, centrifuge briefly
23. Run lig program listed here in step a.
    (a) Choose the preheat lid option and set to 100°C
        i. 30C for 10 minutes
        ii. Hold at 4C
    (b) Add 5 $\mu$L STL to each well, and then mix w/pipette. NOTE- You can hold this mixture at -20C overnight with no trouble.
24. Vortex Sample Purification Beads (SPB) until well dispersed.
25. Perform steps a through l using the Round 1 volumes.
    (a) Add SPB to each well, and then mix thoroughly as follows.
        i.

| | |
|---|---|
| Round 1 | 42.5 $\mu$L |
| Round 2 | 50 $\mu$L |

    (b) Pipette up and down.
    (c) Incubate at room temperature for 5 minutes.
    (d) Place on a magnetic stand and wait until the liquid is clear (2–5 minutes).

(e) Remove and discard all supernatant from each well.

(f) Wash 2 times as follows.

  i. Add 200 $\mu$L freshly prepared 80% EtOH to each well.
  ii. Incubate on the magnetic stand for 30 seconds.
  iii. Remove and discard all supernatant from each well.

(g) Use a 20 $\mu$L pipette to remove residual EtOH from each well.

(h) Air-dry on the magnetic stand for 5 minutes.

(i) Add RSB to each well.

  i.

| | |
|---|---|
| Round 1 | 52.5 $\mu$L |
| Round 2 | 27.5 $\mu$L |

(j) Remove from the magnetic stand, and then mix thoroughly as follows. Pipette up and down.

(k) Incubate at room temperature for 2 minutes.

(l) Place on a magnetic stand and wait until the liquid is clear (2–5 minutes).

(m) CONTINUE TO STEP 26

26. ROUND1- Transfer 50 $\mu$L supernatant to the corresponding well of the CAP plate.

27. Repeat steps a through l with the new plate using the Round 2 volumes.

28. ROUND2- Transfer 25 $\mu$L supernatant to the corresponding well of the PCR plate.

SAFE STOPPING POINT If you are stopping, seal the plate and store at -25C to -15C for up to 7 days.

At this point, ligation and purification is done.

The following is the recipe for each ligated sample:

| Sample | Concentration | Volume ($\mu$L) |
|---|---|---|
| DNA mix | | 3 |
| Forward post-ligation primer (see **Supplemental Section 1**) | 10˘M | 1.5 |
| Reverse post-ligation primer (see **Supplemental Section 1**) | 10˘M | 1.5 |
| Kapa HiFi 2x | 2.5x | 12 |
| Molecular Grade Water | | 12 |
| Total | | 30 |

Follow the following thermocycling protocol with the above 30 $\mu$L mixture:

[noitemsep]Choose the preheat lid option and set to 100C 95C for 3 minutes 8 total cycles of:[noitemsep]

- – 98C for 20 seconds

  - 60C for 15 seconds
  - 72C for 30 seconds
- 72C for 3 min
- Hold at 4C

### 3 Modeling Large Scale Pools

We modeled PCR efficiency as the number of files present in a pool increases. This figure shows how the expected percent of target reads in the solutions changes as we use more primer sequences to specify more files, and when we use different numbers of PCR cycles.

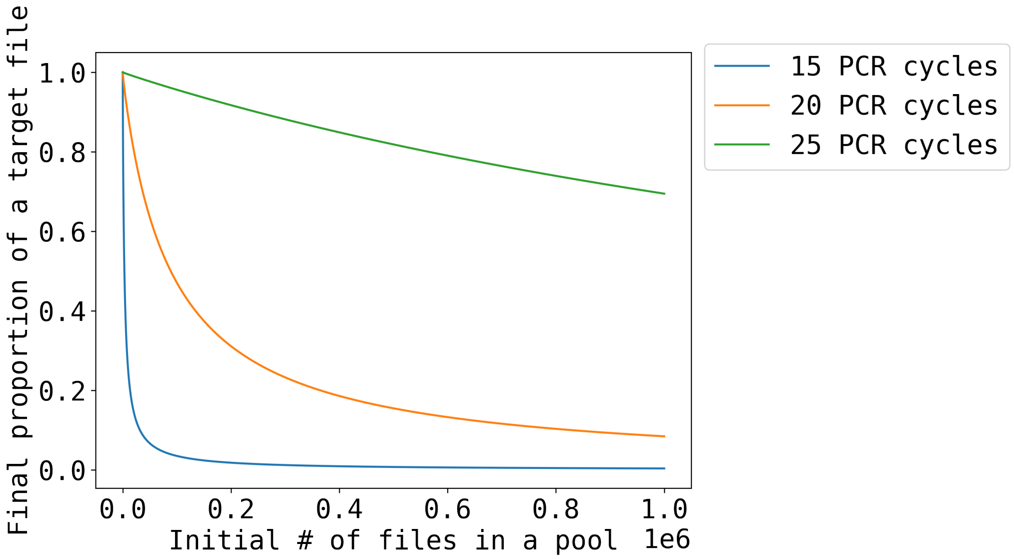


Supplementary Figure 1: Simulation of target file percentage against number of files. Random access efficiency (i.e., proportion of sequences retrieved that correspond to a target file) decreases as more files are stored in a pool. Increasing the number of cycles in a PCR improves the enrichment of a target file, thus increasing the random access efficiency.
